# The Translation Termination Factor eRF3 Interacts Sequentially with eRF1 and ABCE1 on the Ribosome

**DOI:** 10.1101/2025.02.05.636767

**Authors:** Miki Wada, Koichi Ito

## Abstract

To elucidate the molecular mechanisms connecting translation termination and ribosome recycling, interactions among eRF1, eRF3, and ABCE1, were genetically investigated in *Saccharomyces cerevisiae*. Heterologous factors from *Pneumocystis carinii* and humans, along with their mutants were successfully used for co-functional studies in conditional double knockout, double tetOFF strains. Combination of *P. carinii* eRF3 mutants with *S. cerevisiae* eRF1 and ABCE1 revealed that all the eRF3 mutants co-functioned with eRF1, but exhibited varying interactions with ABCE1 depending on the mutants. These findings provide evidence that eRF3 remains bound to the GTPase center of the ribosome, awaiting ABCE1. Thus, eRF3 sequentially interacts with eRF1 and ABCE1 during the final steps of translation.

## Introduction

ABCE1 is a eukaryotic ribosome recycling factor *(1)(2)*, although it was initially discovered as RNase L inhibitor in humans *(3)(4)*. Thus, it is also referred to as Rli1, even in species that lack RNase L such as *Saccharomyces cerevisiae*. ABCE1 consists of an N-terminal FeS cluster domain followed by two structurally and functionally asymmetric ATPase domains *(5)(6)(7)*. ABCE1 remains associated with the 40S ribosomal subunit after the 80S ribosome splitting into subunits *(8)*, and reportedly interacts with translation initiation factors, such as eIF3 subunits *(9)(1)*. Cyro-EM structures capturing eRF1 and ABCE1 on the ribosome suggest that ABCE1 might also play a role in earlier steps of translation, specifically termination *(10)(11)*. While biochemical studies support this idea *(2)(1)*, the precise connection between translation termination and ribosome recycling remains unclear.

In eukaryotes, translation termination is mediated by the release factors eRF1 and eRF3. eRF1 directly recognizes the three stop codons of mRNAs, and cleaves peptidyl-tRNA to release synthesized polypeptide-chain. eRF3, a GTPase dependent on both eRF1 and the ribosome *(12)*, enhances this process by directly binding to eRF1 and functions in concert on the ribosome *(13)(14)(15)*. As many translation termination studies predate the identification of ABCE1 as a ribosome recycling factor, understanding how these two processes are coordinated is crucial. Building on our previous genetic findings *(16)*, this work focuses on clarifying whether eRF3 and ABCE1 interact on the ribosome (Fig. 1A).

**Fig. 1.**
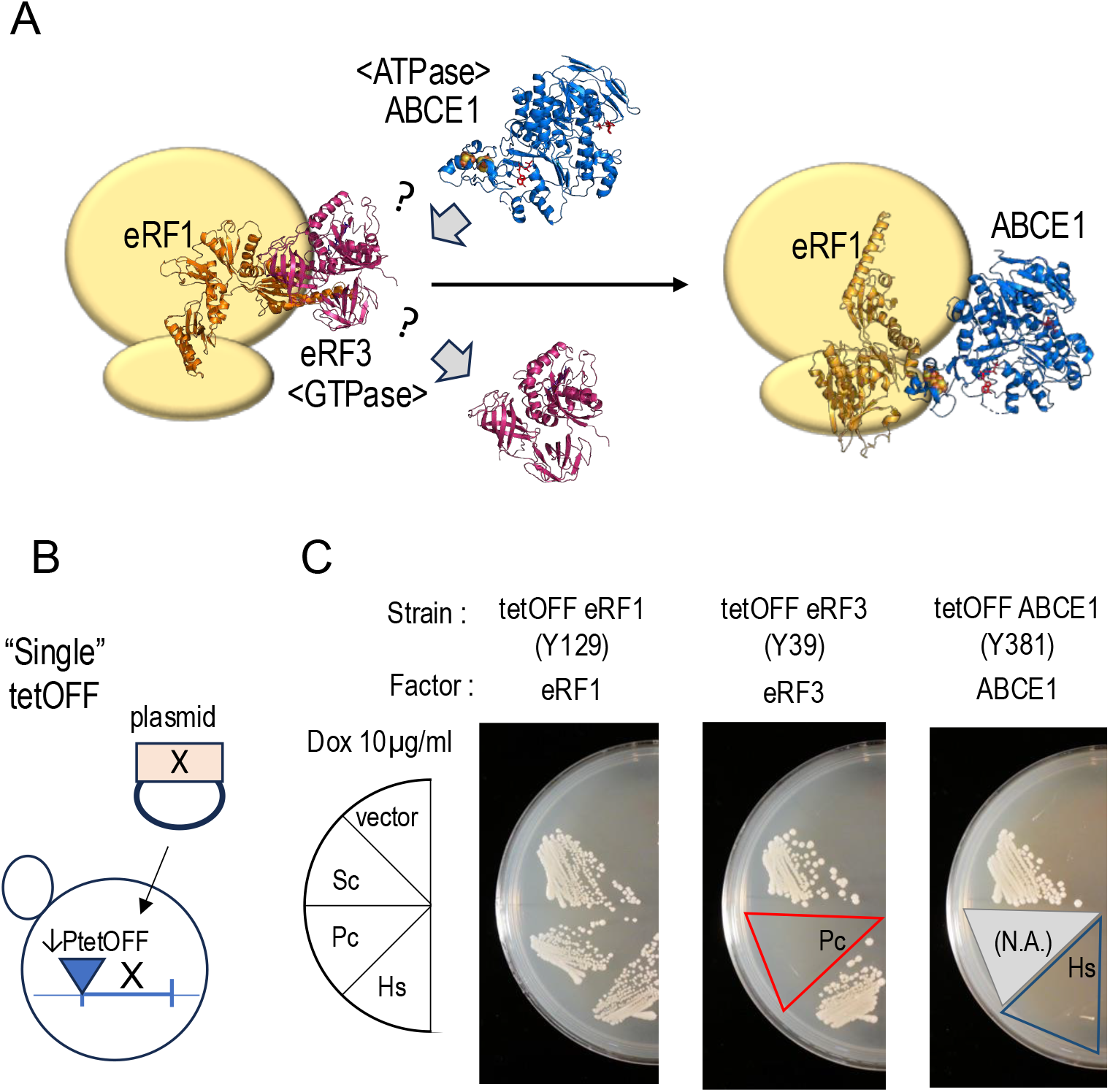
Complementation analysis of tetOFF-eRF1, tetOFF-eRF3, and tetOFF-ABCE1 by corresponding heterologous factors. **A**. Schematic representation of the study’s aim: to determine the order of events during translation termination-does eRF3 dissociate from the ribosome before or after the ribosome recycling factor ABCE1 arrives? **B**. Schematic representation of the complementation analysis. **C**. Complementation of tetOFF-eRF1 strain (Y129) was tested using the p414GPD vector (negative control), and Sc-eRF1 (positive control), Pc-eRF1, and Hs-eRF1 expressed from the vector in the presence of 10µg/ml doxycycline (dox). Complementation of tetOFF-eRF3 (Y39) was tested using the p416GPD vector (negative control), and Sc-eRF3C (positive control), PceRF3C, and s-eRF3a expressed from the vector in the presence of 10µg/ml dox. Complementation of tetOFF-ABCE1 was tested using p415GPD vector (negative control), and Sc-ABCE1(Rli1) (positive control), and Hs-ABCE1 expressed from the vector in the presence of 10µg/ml dox.

## Results

### Complementation of single tetOFF strains by heterologous factors

To assess the functionality of heterologous eRF1s, eRF3s and ABCE1, single tetOFF *S. cerevisiae* strains where the indicated gene expression is repressed by doxycycline (a derivative of tetracycline) were prepared. While some of these factor’s functionality have been individually reported *(16)(17)*, their functionality were examined together in our system. Into tetOFF-eRF1 strain, eRF1 genes from *S. cerevisiae* (Sc-eRF1) (positive control), *P. carinii* (Pc-eRF1), and *Homo sapiens* (Hs-eRF1) were introduced and assessed complementation. Both Pc-eRF1 and Hs-eRF1 showed growth complementation (Fig. 1). In the tetOFF-eRF3 strain, Pc-eRF3C expressed from the encoding plasmid did not complement growth, whereas Hs-eRF3a (encoded by GSPT1) did. In the tetOFF ABCE1 strain, Hs-ABCE1 did not function at 30°C, consistent with our previous report *(17)*. Pc-ABCE1 could not be studied due to the unavailability of the gene and cDNA. *P. carinii* is the causative agent of Pneumocystis pneumonia in immunocompromised host, and is phylogenetically classified as a fungus closely related to *Schizosaccharomyces pombe*. The differential complementation observed with Pc- and Hs-factors prompted us to construct double tetOFF strains for further investigation. Heterologous factors, expected to function optimally in their native species, are likely to malfunction in *S. cerevisiae* due to incompatible interactions with any of *S. cerevisiae* factors.

### Functional interactions among heterologous eRF1, eRF3 and ABCE1

We previously constructed a conditional double-knockout tetOFF-eRF1*(SUP45)*/eRF3*(SUP35)* strain to demonstrate functional interaction between of eRF1and eRF3 *(16)*. To further investigate the functional relationship among these factors, we constructed two new conditional double-knockout strains: tetOFF-eRF3/ABCE1 (Y379) and tetOFF-eRF1/ABCE1 (Y381). Using these three strains, the functional interaction among eRF1, eRF3 and ABCE1 from *S. cerevisiae, H. sapiens*, and *P. carinii* was studied on a two-factor basis (Fig. 2A), while acknowledging crucial role of the ribosome together.

**Fig. 2.**
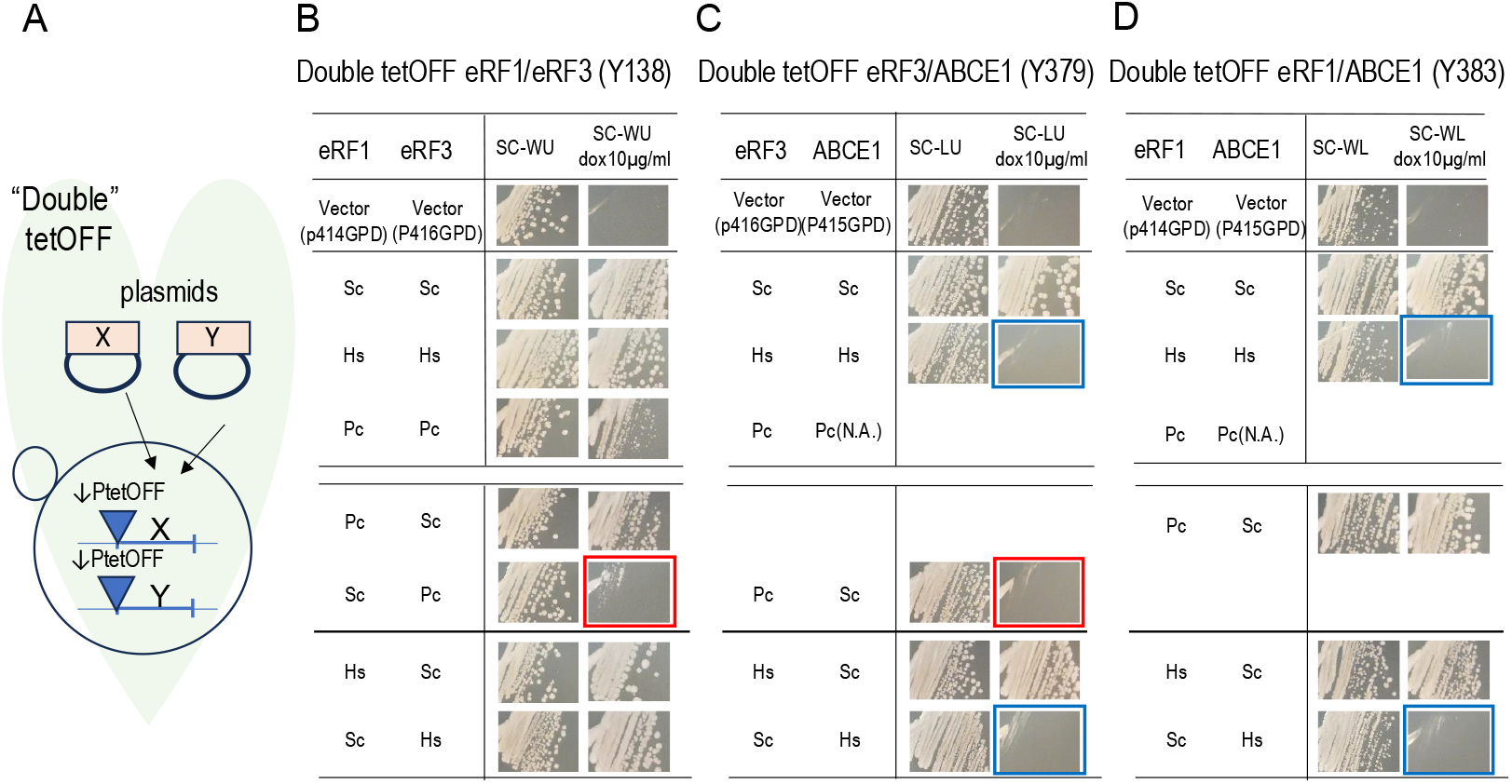
Co-functionality analyses in double tetOFF strains using pairs of eRF1, eRF3, and ABCE1 from *S. cerevisiae, P. carinii*, and *H. sapiens*. **A**. Schematic diagram of the co-functionality analysis in double tetOFF strains. **B**. The indicated combinations (left column) of eRF1 and eRF3 encoding plasmids were introduced into the double tetOFF-eRF1/eRF3 strain, and growth was examined with and without 10µg/ml dox (right column). **C**. The indicated combinations of eRF3 and ABCE1 encoding plasmids were introduced into the double tetOFF-eRF3/ABCE1 strain, and growth was examined with and without 10µg/ml dox. **D**. The indicated combinations of eRF1 and ABCE1 encoding plasmids were introduced into the double tetOFF-eRF1/ABCE1 strain, and growth was examined with and without 10µg/ml dox. Throughout Fig. 2, factors from *S. cerevisiae* are expressed from ADH promoter vector, while factors from *P. carinii* and *H. sapiens* are expressed from a GPD promoter vector.

Expression of pairs of corresponding factors from *S. cerevisiae*, Sc-eRF1, Sc-eRF3C, and Sc-ABCE1, in each strain supported growth under 10 µg/ml doxycycline (dox) conditions, confirming the genotypes of these strains (Fig. 2B-D, Sc-Sc combinations). We used C-terminal region of eRF3 (eRF3C), responsible for translation termination, to avoid potential overproduction effects. In the tetOFF-eRF1/eRF3 strain, expression of two corresponding factors from the same species, either *H. sapiens* or *P. carinii*, allowed growth (Fig. 2B, Hs-Hs, Pc-Pc). However, the Pc-eRF1/Pc-eRF3C combination showed reduced growth, suggesting that beyond this two-factor interaction might influence the phonotype. In the tetOFF-eRF3/ABCE1 and tetOFF-eRF1/ABCE1 strains, combinations of human factors did not support growth (Fig. 2C and 2D, Hs-Hs combinations), likely due to functional incompatibility between Hs-ABCE1 and *S. cerevisiae* machinery, most likely the ribosome. Hs-ABCE1 did not show functionality in any of the examined combinations (Fig. 2C-D, blue boxes), and co-functionality with Pc-ABCE1 could not be assessed due to its unavailability.

In two-factor-hetero combinations, Pc-eRF1 functioned with both Sc-eRF3C and Sc-ABCE1. However, as observed previously *(16)*, Pc-eRF3C could not function with Sc-eRF1 (Fig. 2B, red box). Importantly, Pc-eRF3C also failed to function with Sc-ABCE1 (Fig. 2C, red box). In summary, Pc-eRF3 cannot function with either Sc-eRF1 or Sc-ABCE1 in the double knockout strains, suggesting that eRF3 interacts with both eRF1 and ABCE1. We further investigated these interactions using these combinations as a tool.

### Dual co-function of Pc-eRF3C mutants

We previously isolated Pc-eRF3C mutants that complemented the growth of either temperature-sensitive *sup35* strain or tetOFF-eRF3 strains *(16)*. These mutants were used for the hetero-compatibility study shown in Fig. 3A. All twelve Pc-eRF3C mutants exhibited co-functionality with Sc-eRF1 in double tetOFF-eRF1/eRF3 strain (Fig. 3B-C, left). Next, we examined the co-functionality of these mutants with Sc-ABCE1 in the double tetOFF-eRF3/ABCE1 strain (Fig. 3B-C, right). Intriguingly, the Pc-eRF3C mutants displayed differential functionality with Sc-ABCE1: six out of 12 showed functionality (Fig. 3C, counting + and ++), although some weakly. In summary, six Pc-eRF3C mutants functioned with both Sc-eRF1 and Sc-ABCE1. This result provides evidence for a direct interaction between eRF3 and ABCE1 on the ribosome. If eRF3 dissociated from the ribosome before ABCE1’s arrival, these eRF3 mutants would not be expected to show differential functionality with ABCE1, suggesting a novel model of translation termination leading to ribosome recycling. The locations of Pc-eRF3C mutants that co-function with ABCE1(+ or ++ in Fig. 3C) are on the GTP-binding side of eRF3, suggesting a potential interaction surface with ABCE1 (Fig. 3D). While not all positive mutations necessarily correspond to direct contact sites, the positions of these mutations on the eRF3 3D structure suggest that at least some may be involved in direct contact with Sc-eRF1 and Sc-ABCE1.

**Fig. 3.**
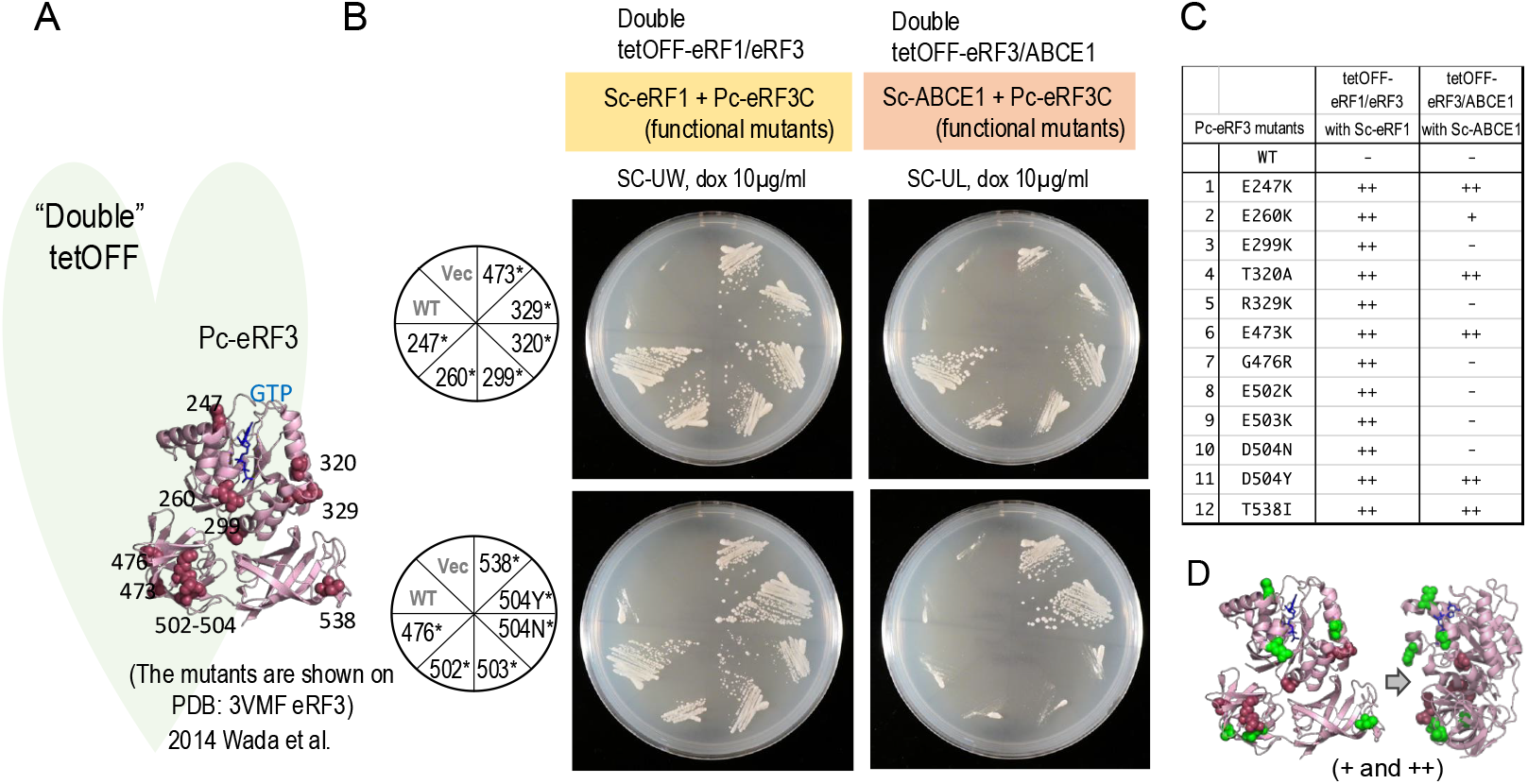
Co-functionality of Pc-eRF3C and its mutants with Sc-eRF1 and Sc-ABCE1. **A**. Schematic of co-functionality analyses in the double strains, with the locations of Pc-eRF3 mutations mapped onto the protein structure. **B**. (Left) Plasmids encoding wild-type Pc-eRF3C and its mutants, along with Sc-eRF1, were introduced into the double tetOFF eRF1/eRF3 strain, and growth was assessed on plates containing 10µg/ml dox. (Right) Plasmids encoding wild-type Pc-eRF3C and its mutants, along with Sc-ABCE1, were introduced into the double tetOFF eRF3/ABCE1 strain, and growth was assessed on plates containing 10µg/ml dox. Sc-eRF1 was expressed from a p414ADH vector, Sc-ABCE1 from a p415ADH vector, and Pc-eRF3C and its mutants from a p416GPD vector. **C**. Summary of growth results from panel **B**, indicating the co-functionality of Pc-eRF3C mutants with Sc-eRF1 and Sc-ABCE1. **D**. Schematic representation of co-functional Pc-eRF3C mutants with Sc-ABCE1 (+ and ++ in panel C right) in green. Throughout Fig. 3, Pc-eRF3C and its mutants were expressed from a GPD promoter, while Sc-eRF1 and Sc-ABCE1 were expressed from a medium-expression ADH promoter.

### Loss-of-function analysis of the Sc-eRF3 E3GGQ motif

The evidence for a direct interaction between the GTPase eRF3 and the ATPase ABCE1, presumably involving the nucleotide-binding side of eRF3, prompted a careful investigation of the nucleotide-binding motifs of both molecules (Fig. 4A). As a translational GTPase, eRF3 contains conserved G1 to G4 motifs *(18)(19)*, shared with elongation GTPases. Intriguingly, semi-conserved motif between G3 and G4 differs between eRF3 and other translational GTPases: “GGQTREH” in eRF3s versus “xxQTREH” (where x is any amino acid) in other translational GTPases. This “GGQ” is conserved exclusively in eRF3s and is designated here as the eRF3-specific GGQ (E3GGQ) motif (Fig. 4A). The reason we focused on the motif is that, in ABC (ATP binding cassette) proteins, LSGGQ is well-known signature motif involved in interaction with gamma-phosphate of the nucleotide *(20)(21)*, though it is LSGGE in ABCE1. The conservation of “GGQ” in both eRF3 and ABC proteins is intriguing, given their potential interaction with gamma-phosphate. The E3GGQ motif might affect co-function with eRF1, ABCE1, or both, potentially via GTP nucleotide related interaction. To investigate this, site-directed E3GGQ motif mutants of Sc-eRF3 were constructed and examined co-functionality in homologous *S. cerevisiae* system to assess loss-of-function effect (Fig. 4B).

**Fig. 4.**
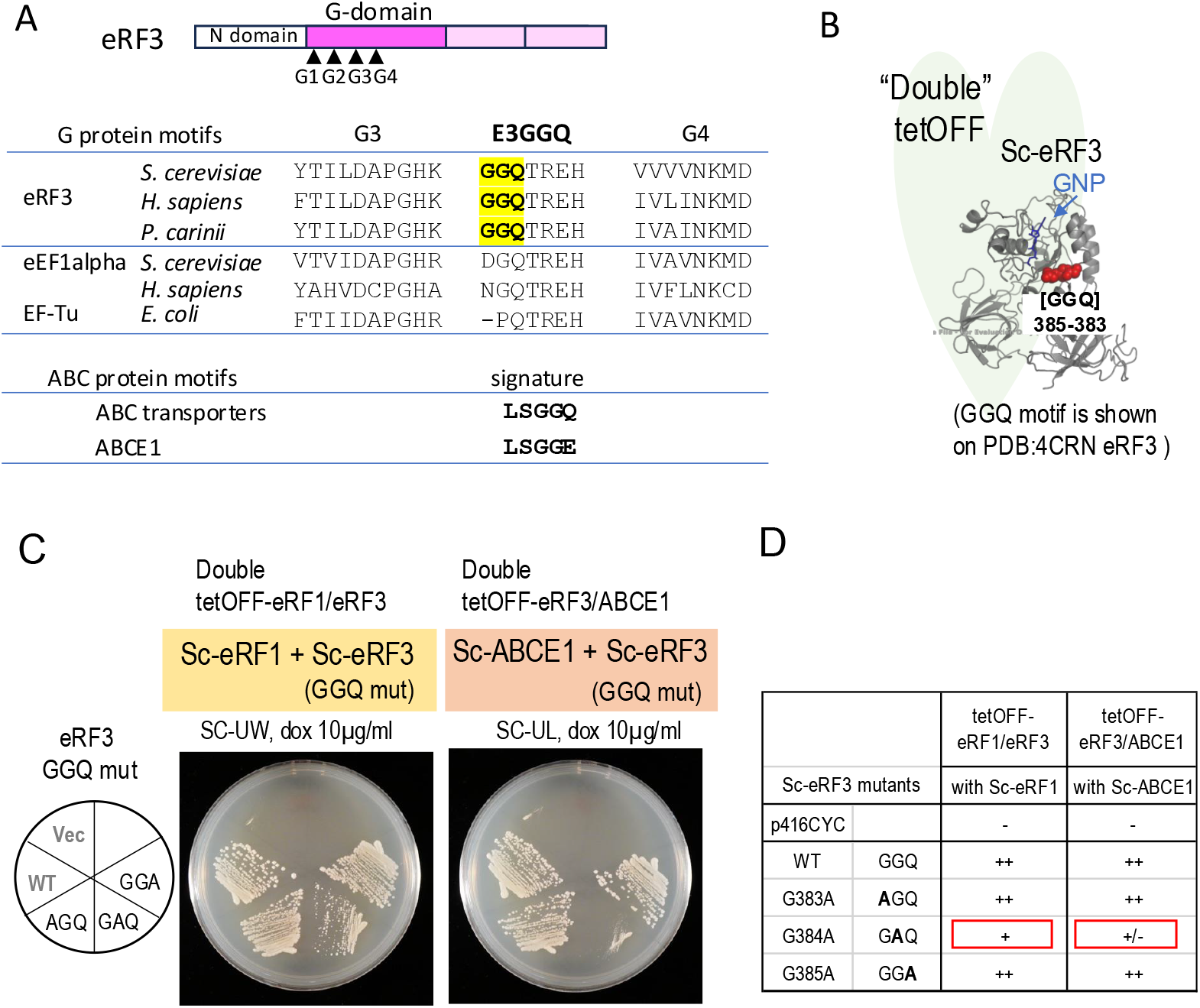
Effect of Sc-eRF3C GGQ motif mutations on co-functionality with homologous Sc-eRF1 and Sc-ABCE1 factors. **A**. The G-protein motifs, G3 and G4, of representative species are shown aligned. A newly identified eRF3-specific GGQ motif, designated E3GGQ, is highlighted in yellow background. The related, well-known ABC protein “signature motif” is shown for comparison. **B**. Schematic of the co-functionality analysis, highlighting the location of the Sc-eRF3 GGQ motif in the structure. **C**. (Left)Plasmids encoding the Sc-eRF3C and its E3GGQ motif mutants (G383A, G384A, and G385A), along with Sc-eRF1, were introduced into the double tetOFF-eRF1/eRF3 strain, and growth was assessed on plates containing 10µg/ml dox. (Right) Plasmids encoding the Sc-eRF3C and its E3GGQ motif mutants (G383A, G384A, and G385A), along with Sc-ABCE1, were introduced into the double tetOFF-eRF3/ABCE1 strain, and growth was assessed on plates containing 10µg/ml dox. **D**. Summary of the growth results from panel **C**, indicating the co-functionality of eRF3 GGQ motif mutants with either Sc-eRF1 or Sc-ABCE1. All factors in Fig.4 are expressed from low-expression CYC1 promoter.

Each amino acid of GGQ from the Sc-eRF3 E3GGQ motif (residues 383-385) was mutated to alanine, and these Sc-eRF3C mutants were tested for their co-functionality with both Sc-eRF1 and Sc-ABCE1 (Fig. 4C-D). One mutant GAQ (G384A), exhibited only a minor growth defect compared to wild-type eRF3C in combination with Sc-eRF1 in double tetOFF-eRF1/eRF3 strain (Fig 4C-D, left). However, in the tetOFF-eRF3/ABCE1 strain, the GAQ mutant showed significant growth reduction with wild-type ABCE1 (Fig. 4C-D, right). Thus, the GAQ eRF3 mutant has a more pronounced effect on the interaction with ABCE1 than with eRF1, suggesting that the GTPase eRF3 and the ATPase ABCE1 interact via a GGQ motif-dependent mechanism. This confirms an interaction between eRF3 and ABCE1 in a homologous *S. cerevisiae* system (Fig. 4. Fig. 5).

**Fig. 5.**
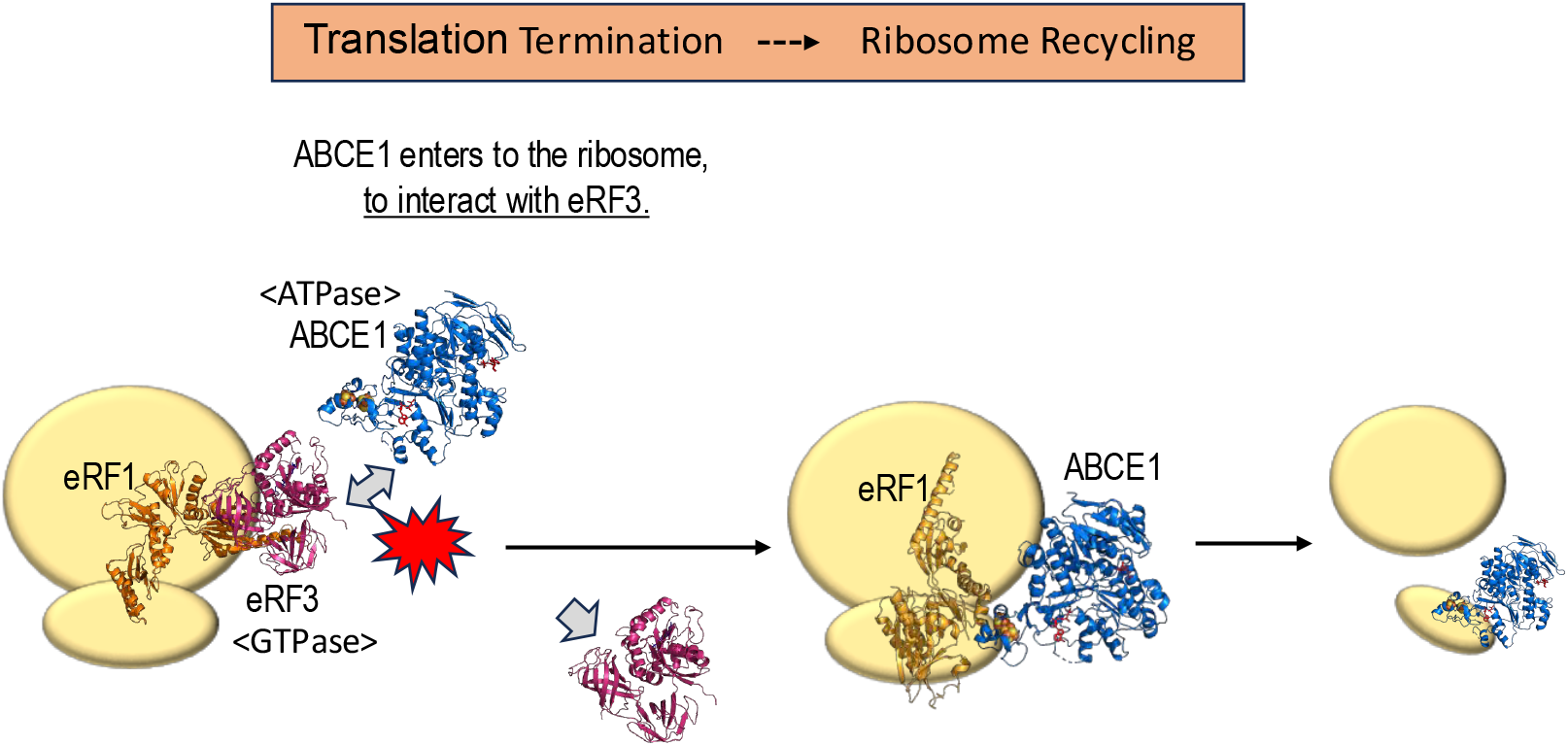
Model illustrating the sequential interaction between eRF1 and eRF3, and ABCE1 on the ribosome. After translation termination mediated by eRF1 and eRF3 through its GTPase function, eRF3 remains on the ribosome, awaiting interaction with ABCE1. This interaction triggers the next step, accompanied by conformational changes, leading to ribosome recycling.

## Discussion

The non-functional heterologous factor Pc-eRF3 was proved to be a valuable tool for demonstrating molecular interactions on the ribosome. Using double tetOFF strains, we surprisingly fond that Pc-eRF3 is incompatible with both Sc-eRF1 and Sc-ABCE1 in two-factor basis studies (Fig. 2). While the ribosome and other factors should undoubtedly play a role, the lack of growth in these experiments appears to be primarily due to a direct incompatible interaction between Pc-eRF3 and these two factors on the ribosome. The use of double tetOFF strains was instrumental in elucidating these interactions. Applying this double tetOFF strain approach to other suitable systems could significantly enhance our understanding of molecular interactions, particularly within complex macromolecular assemblies.

Further analyses with Pc-eRF3C growth-complementing mutants revealed that all 12 Pc-eRF3 mutants co-functioned with Sc-eRF1, suggesting that eRF1/eRF3 co-functionality is a prerequisite for the subsequent steps. In co-functionality experiments between Pc-eRF3 mutants and Sc-ABCE1, differential compatibility was observed depending on the specific mutations (Fig. 3). This provides evidence for a direct interaction between Pc-eRF3C and Sc-ABCE1. If eRF3 were to dissociate from the ribosome before ABCE1’s arrival, the eRF3 mutants would not be expected to affect the subsequent interaction with ABCE1. Therefore, our data support a model where eRF3 interacts sequentially with eRF1 and then ABCE1 on the ribosome (Fig. 5).

The finding that the interaction between eRF3 and ABCE1 raises many intriguing questions warrants further investigation. eRF3 is a GTPase dependent on eRF1 and the ribosome, which enhances translation termination. In related 3D structures, eRF3 and ABCE1 occupy similar site on the ribosome *(10)(11)*. Translational GTPases, such as eEF1A, eEF2 and eRF3 in eukaryotes, interact with so-called GTPase associated center (GAC) on the ribosome through conserved interaction motifs. However, ABCE1 interacts with the ribosome in a completely different manner as a non-GTPase molecule. Given that co-functional mutation sites of Pc-eRF3 interact with ABCE1, it is supposed that the interaction occurs in a face-to-face manner, with their nucleotide-binding faces directly interacting. This raises an interesting working model in which the gamma-phosphate can be exchanged between GTP/GDP bound eRF3 and ATP/ADP bound ABCE1. The translation elongation GTPases, eEF1A in eukaryotes and EF-Tu in prokaryotes, use guanine nucleotide exchange factor (GEF) to exchange their binding GDP for GTP, along with a conformational exchange between GDP-bound and GTP-bound states *(22)(23)(24)*. Although eRF3 shares similarity with eRF1A, GEF for eRF3 has not been identified. It is possible that ABCE1 could function as a GEF-like protein for eRF3 in a novel mechanism. ABCE1 has two ATP-binding sites that are reported to function asymmetrically (6)(7). One of these ATP-binding sites might facilitate the exchange of phosphate groups with the nucleotide bound to eRF3.

“GGQ” motif is well studied in eRF1, where it is locates at the tip of domain 2 and is presented to the peptidyl transferase center (PTC) of the ribosome to facilitate peptidyl-tRNA hydrolysis *(25)(26)*. Interestingly, ATP-binding cassette (ABC) transporter superfamily proteins share a related motif, the “signature motif” (“LSGGQ”), and ABCE1 has similar motif, “LSGGE”. The “LSGGQ” /”LSGGE” motifs have been reported to interact with the gamma-phosphate group of ATP molecule *(20)(27)*. Furthermore, eRF3 has a conserved “GGQ” motif in close proximity to the gamma-phosphate of GTP (Fig. 4A), although this has not been reported in previous studies to the best of our knowledge. The motif is absent in homologous translational GTPases such as eEF1A, EF-Tu, or aEF1A (archaeal EF1A, which functions as both aEF1A and aRF3) *(28)*, but is found exclusively in eRF3. Co-functionality experiments with the E3GGQ motif mutants (Fig. 4), revealed that one of the mutants, GAQ, plays an important role in co-functioning with ABCE1. These results support our working model of face-to-face interaction between eRF3 and ABCE1, potentially involving their nucleotide-binding sites. The conformational changes of eRF1, eRF3 and ABCE1, accompanied with GTPase and ATPase reactions, likely drive the transition from translation termination to ribosome recycling. There may be transient conformations that have yet to be uncovered. Our genetic study has captured a transient molecular interaction between eRF3 and ABCE1 that would be difficult to study using other methods.

## Acknowledgments

I (M. Wada) thank the Graduate School of Frontier Sciences (GSFS), University of Tokyo, for treating me as a guest scientist.

## Author contributions

Conceptualization, Methodology, Investigation, Visualization: M.W. Funding acquisition: K.I. Project administration: M.W. Supervision: M.W., K.I. Writing – original draft: M.W. Writing – review & editing: M.W., K.I.

## Data and materials availability

All data are available in the main text or the supplementary materials. *P. carinii* eRF1 and eRF3 cDNA sequences were deposited to the database in the following numbers, DDBJ/EMBL/GenBank AB052893 and AB052894 (29).

## Materials and Methods

### Yeast media and strains

Synthetic complete media was used for yeast plates prepared with indicated dropout mix (Formedium, Swaffham, UK), supplemented with 10µg/ml doxycycline (dox) where indicated. Experimental procedures for *S. cerevisiae* are conventional ones. *S. cerevisiae* transformations were conducted with Frozen-EZ transformation II kit (Zymoresearch).

The strains used are the following.

tetOFF-eRF1 (Y129): MAT*α* his3Δ200 leu2Δ0 lys2Δ0 met15Δ0 trp1Δ63 ura3Δ0 tetOFF(hphMX-tTA-tetO)-*SUP45*

tetOFF-eRF3 (Y39): MAT*α* ade2::hisG his3Δ200 leu2Δ0 lys2Δ0 met15Δ0 trp1Δ63 ura3Δ0 HO-tetOFF(hphMX-tTA-tetO) -*SUP35*-HO sup35::HIS3

tetOFF-ABCE1 (Y381): MAT*α* his3Δ200 leu2Δ0 lys2Δ0 met15Δ0 trp1Δ63 ura3Δ0 tetOFF(kanMX-tTA-tetO) -*Rli1*

double tetOFF-eRF1/eRF3 (Y138): MAT*α* ade2::hisG his3Δ200 leu2Δ0 lys2Δ0 met15Δ0 trp1Δ63 ura3Δ0 HO-tetOFF(kanMX-tTA-tetO) -*SUP35*-HO sup35::HIS3 tetOFF(hphMX-tTA-tetO)-*SUP45*

Double tetOFF eRF3/ABCE1 (Y379): MAT*α* ade2::hisG his3Δ200 leu2Δ0 lys2Δ0 met15Δ0 trp1Δ63 ura3Δ0 HO-TetOFF(hphMX-tTA-tetO) -*SUP35*-HO sup35::HIS3 tetOFF(kanMX-tTA-tetO)-*Rli1*

Double tetOFF eRF1/ABCE1 (Y383): MAT*α* his3Δ200 leu2Δ0 lys2Δ0 met15Δ0 trp1Δ63 ura3Δ0 tetOFF(hphMX-tTA-tetO)-*SUP45* tetOFF(kanMX-tTA-tetO)-*Rli1*

### Yeast plasmids

The plasmids used in this study are listed in the following table.

The cDNA sequences of Pc-eRF1 and Pc-eRF3 are posted in DDBJ/EMBL/Genbank AB052893 and AB052894, respectively *(1)*.

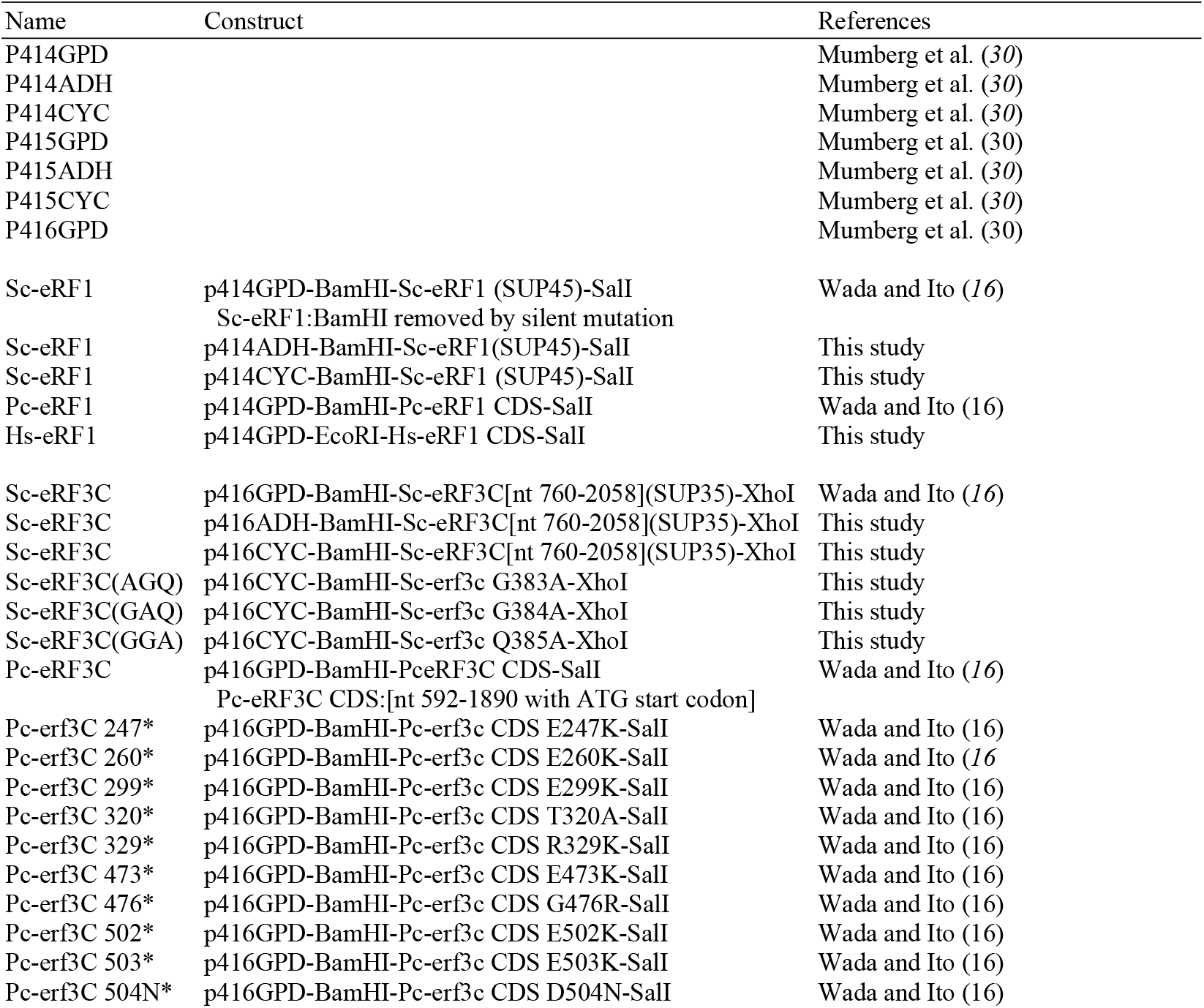

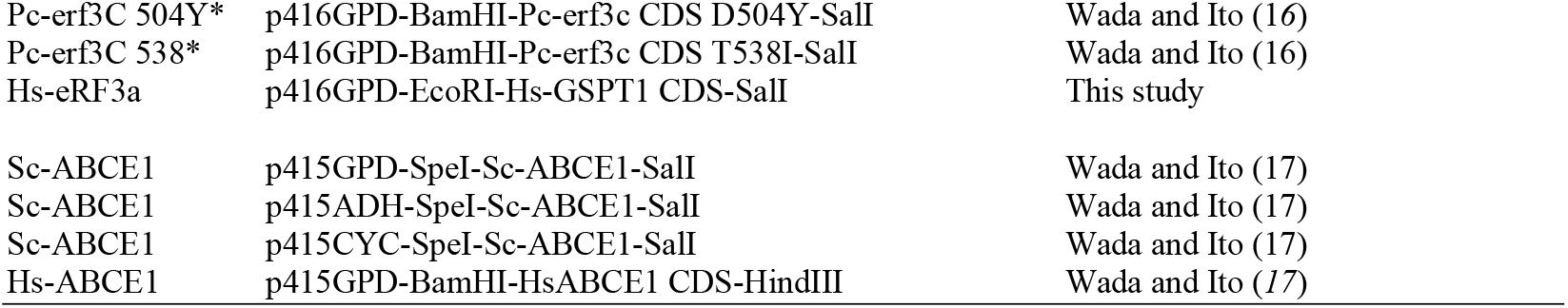

### Site directed mutagenesis

The GGQ mutations were introduced by 2 step PCR with the following mutagenesis primers. The 1^st^ PCR was performed from wild type Sc-eRF3 encoded plasmid, by combinations of promoter primer and mutagenesis primer1, and CYC terminator primer and primer2. The 2^nd^ PCR was performed from the two fragments of 1^st^ PCR with promoter primer and CYC terminator primer. The obtained fragments were cloned into the p416CYC vector.

AGQ: primer1a; TTCACGAGTTTGACCAGCTCTCTCAAAACCGGT, primer2a; ACCGGTTTTGAGAGAGCTGGTCAAACTCGTGAA

GAQ: primer1b; GTGTTCACGAGTTTGAGCACCTCTCTCAAAACC, primer2b; GGTTTTGAGAGAGGTGCTCAAACTCGTGAACAC

GGA: primer1c; GGCGTGTTCACGAGTTGCACCACCTCTCTCAAA, primer2c; TTTGAGAGAGGTGGTGCAACTCGTGAACACGCC

